# Active locomotion predictively rescues head direction attractor dynamics in head-fixed mice

**DOI:** 10.64898/2026.01.12.698940

**Authors:** Alexandr Pak, Janna Aarse, Heng Wei Zhu, Jean-Paul Noel, Simón Carrillo Segura, André A. Fenton, Dora Angelaki

## Abstract

Head direction (HD) cells in the anterodorsal thalamic nuclei form the brain’s internal compass, and are often modeled as a ring attractor maintaining azimuth coding by leveraging continuous visual and inertial sensory input. Here, we test how the common experimental preparation of head-fixed animals alters this code. Complete head-fixation that creates vestibular conflict disrupts both unit and population encoding of head direction, while selectively constraining head-on-body movements either in real or virtual reality uniquely impairs HD population activity. More specifically, attractor dynamics is altered in head-restrained mice during periods of immobility, but remarkably recover several hundred milliseconds prior to locomotion onset. The rescue preceding movement onset suggests that an efference copy or prediction of a re-afferent signal is necessary to maintain HD network activity during head restraint. A computational model recapitulates these effects by perturbing lateral connectivity among HD neurons. More generally, the results indicate that the HD network is a context- and state-dependent predictive estimator, stabilized by forthcoming self-motion signals. The classic ring-attractor models should be revised to integrate context-dependent dynamics with prospective motor signals, offering a more complete account of how the brain’s compass remains stable across both naturalistic and constrained conditions.

## Introduction

The mammalian navigation system relies on specialized neural ensembles that encode spatial variables critical for orientation and wayfinding^1,2^. Among these, head direction (HD) cells in the anterodorsal thalamic nucleus provide a compass-like signal that supports higher-order spatial representations throughout the brain, as evidenced by the abolition of grid cell firing patterns in the medial entorhinal cortex following thalamic inactivation^3,4^. The anterior thalamic HD signal is classically conceptualized as a one-dimensional ring attractor network encoding azimuthal orientation^5–7^, where neurons maintain consistent directional tuning and network stability through recurrent connectivity.

Recent discoveries have unveiled unexpected computational complexity in the HD system. Beyond simple directional coding, HD cells exhibit three-dimensional tuning to both azimuth and gravitational tilt^8,9^, and implement gain modulation during uncertainty^10^. Moreover, the HD system demonstrates remarkable internal dynamics independent of immediate sensory input. HD cells can maintain stable representations through internally generated activity during sleep via sharp wave ripples^11^, and MEC HD cells can create a stable, internal compass—an egocentric “sense of direction” independent of external landmarks^12^. These findings challenge the passive sensory framework, revealing that HD cells operate through complex multimodal integration and exhibit remarkable behavioral state dependency.

The HD system’s dependence on multimodal integration becomes particularly relevant when considering the increasing prevalence of head-fixed preparations in neuroscience. These paradigms, while enabling advanced techniques like two-photon imaging and precise virtual reality control^13,14^, fundamentally disrupt the natural sensory-motor contingencies that spatial circuits evolved to process. Head fixation may also profoundly affect behavioral state, which influences neural processing throughout the brain, with locomotion known to modulate sensory processing, attention, and neuromodulatory tone^15–17^. Critically, head restraint disrupts normal HD cell function: individual HD cells show reduced directional modulation—altered firing at both preferred and non-preferred directions—during head fixation^18^, suggesting that natural movement is essential for canonical HD network dynamics. Despite the increasing prevalence of head-fixed approaches in navigation research, their specific effects on HD circuit dynamics and potential interactions with behavioral state remain inadequately characterized.

This experimental context raises critical questions: Does head fixation fundamentally alter the computational properties of the HD system? How does behavioral state (immobility versus locomotion) interact with head fixation to influence HD network dynamics? Answering these questions is essential not only for interpreting the extensive neuroscience literature using head-fixed preparations, but also for understanding the fundamental principles governing spatial representation. To address these questions, we developed an experimental approach combining chronic high-density silicon probe recordings across four conditions: freely moving mice, head-restrained yet freely moving mice, yaw-enabled VR, and completely head-fixed VR. The combination of these setups allowed us to isolate the specific effects of head fixation, motor actions, and motor predictions on HD network neural dynamics.

We discovered that the HD system operates as a state-dependent predictive circuit rather than a passive sensory integrator. Head fixation induces an “off-ring” manifold state where population activity moves toward the center of the manifold, driven by reduced directional modulation of HD cells. This “off-ring” state of the HD network is driven by immobilization of head, and emerges selectively during stationary periods. Remarkably, locomotion alone—without head rotation, angular velocity, or translational vestibular input—restores canonical ring dynamics. This rescue of population dynamics occurs on average ∼550ms prior to locomotion onset, suggesting it is driven by motor efference or predictions of re-afference. These findings reveal that motor efference/re-afference can maintain spatial representations independent of sensory feedback, fundamentally revising our understanding of how the brain’s compass operates under both naturalistic and constrained conditions.

## Results

### Active yaw rotations are required for head direction cell tuning in 2D virtual reality

We performed chronic extra-cellular single-cell recordings (Neuropixel 2.0) in the anterodorsal thalamic nucleus (ADN) of six mice (7 implantations, n = 28 sessions) as they engaged in foraging behaviors in two environments. The first environment was a traditional, real open field arena, where the environmental lights may be on (*Real Light*; RL; **Fig. 1A**, orange) allowing the animal to perceive landmarks, or they could be off (*Real Dark*; RD; **Fig. 1A**, gray) requiring animals to navigate via path integration and cues on the apparatus substrate. Under both lights on and off conditions, the animal has access to yaw, pitch, and roll signals. The second environment was a two-dimensional (2D) virtual reality (VR) environment. There were also two VR conditions: First, with the head entirely fixed. Second, mice were allowed to freely rotate in yaw on a bearing while head-fixed (**Fig. 1B**; see^14,19^ for a similar approach). The first VR condition (*VR-YawFix*) induced vestibular conflict. The second VR condition (*VR-YawFree*) eliminated that conflict by allowing for yaw (but not pitch or roll) rotation signals. The VR experiments immediately followed the recordings in the real arena, enabling us to track the same cells across different environments. To motivate mice to navigate in 2D VR they were trained in a target-pursuit task (**Figure S1**).

**Figure 1.**
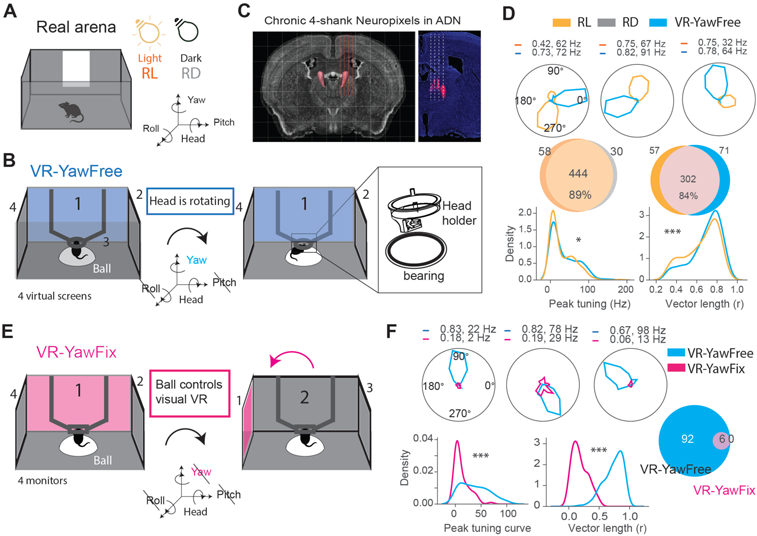
Active head yaw rotations are required for head direction cell tuning in 2D virtual reality. **(A)** Freely moving mice explore an open-field arena under both light and dark conditions. **(B)** Novel 2D virtual reality setup featuring a custom bearing system that permits mice to actively yaw rotate their heads while pitch and roll remain fixed. **(C)** Histology showing a 4-shank Neuropixel 2.0 probe chronically implanted into the anterodorsal thalamic nucleus (ADN). **(D)** Top: Example HD cell tuning curves for real arena during light (RL, orange) and VR-YawFree (blue), and their corresponding Rayleigh vector lengths and peak tuning curve amplitudes. Middle: Pie chart of the number of HD cells in RL vs RD (left) and RL vs VR-YawFree (right). Bottom: Distribution of peak tuning curve amplitude (left) and Rayleigh vector length (right). **(E)** Conventional VR setup in which the mouse’s head remains completely fixed (VR-YawFix) while the visual environment rotation is coupled to the ball rotation, eliminating all vestibular rotation cues. (**F**) Same as (**D**) but comparing VR-YawFree and VR-YawFix. *p < 0.05, **p < 0.01, ***p < 0.001.

We recorded 359 head direction cells (HDCs; 11-40 simultaneously per session) in the ADN (see **Fig. 1C** for example histology) in freely moving mice during RL sessions. The vast majority of these cells (84%) were also identified as HDCs in the VR-YawFree recording, suggesting that active head yaw rotation is sufficient to maintain thalamic HD tuning in the virtual environment (**Fig. 1D**, as a comparison, 89% were maintained between RL and RD), as previously reported for MEC HD cells^14,20^. In fact, both peak firing rates (Mann-Whitney U test, p = 0.02) and Rayleigh vector lengths (p = 5.53e-3) of heading tuning functions were larger in VR-YawFree than RL.

To dissect the role of active yaw vestibular cues in shaping the HD network within VR, we clamped the head-fixating bearing in a subset of sessions to introduce vestibular conflict (**Fig. 1E**, *VR-YawFix*) while still allowing the animals to (visually) forage. Without any vestibular cues, HD cells lost their “directional” tuning and firing patterns became broadly tuned (**Fig. 1F**, VR-YawFree vs VR-YawFix peak tuning amplitude: p = 2.87e-9; Rayleigh vector length, p = 8.77e-26). These results align with earlier reports of the loss of spatial tuning in a traditional 2D VR environment without yaw rotations^13^. Together, these results show that yaw rotation cues are required for proper HD cell tuning in VR. In the presence of visual/vestibular conflict, HDC tuning is lost or severely compromised.

Critically, the HD system is inherently defined by pairwise co-firing patterns rather than single cell coding alone, making population analyses essential for understanding system-level function. Next we examine pairwise coupling and population responses in the stimulus conditions without vestibular conflict by comparing VR-YawFree and RL/RD conditions.

### Head fixation alters population dynamics of head direction networks

Having confirmed that single HD cells maintain proper directional tuning in VR while animals are allowed to yaw rotate (VR-YawFree), we next examined the strength of pairwise correlations between HD cell pairs (mean τ corr) and assessed whether these relationships were preserved across real and VR environments (Δ1). Specifically, we computed Kendall’s tau correlations between the firing rates of HD cell pairs within each environment and then quantified Δτ as the difference in correlation strength for the same cell pairs across the two environments.

This analysis revealed similar pairwise correlation structure in RL and RD sessions (**Fig. 2A**, top and **Fig. 2B**, left for an example session), but weaker pairwise correlation within the VR-YawFree environment than either of the real-world environments (**Fig. 2A**, bottom; **Fig. 2B**, right; example session). Indeed, on average we found significantly weaker pairwise correlations in the VR-YawFree compared to both RL (**Fig. 2C** mean τ corr, p = 0.008) and RD (**Fig. 2C** mean τ corr, p = 0.01), with no difference across the latter two conditions (p = 0.8). Furthermore, while HD cell pair correlations were similar between real environments (RL-correlated pairs remained correlated in RD), they showed extensive remapping in VR-YawFree, where strongly correlated real-environment pairs often became weakly correlated, or anti-correlated (**Fig. 2D** abs (Δτ), p = 1.1e-6). These findings indicate that while single cell tuning is still present, head fixation in VR-YawFree alters co-firing properties of HD cells.

**Figure 2:**
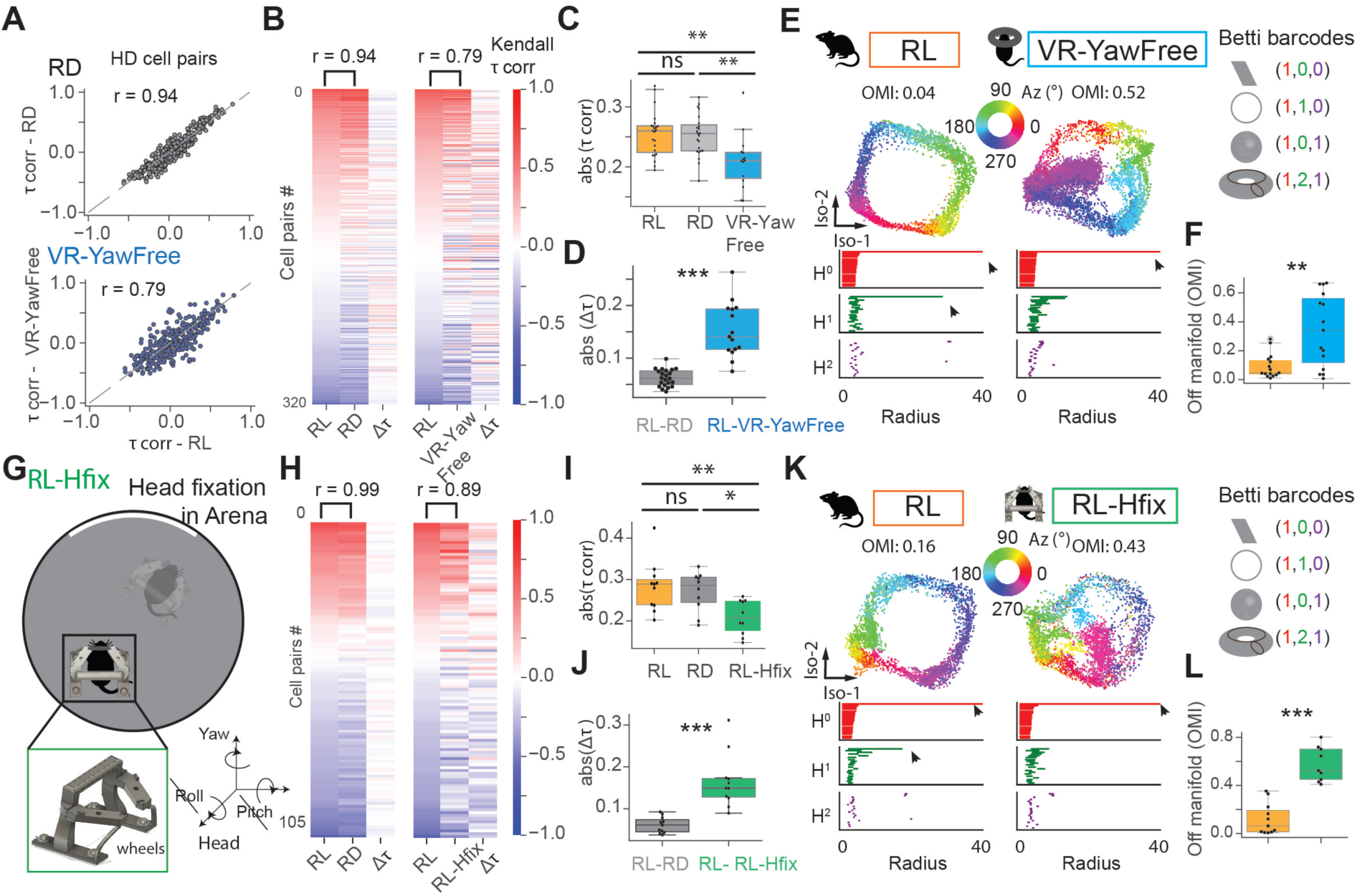
Head fixation alters head direction ring manifold dynamics. **(A)** Scatter plot of pairwise Kendall τ correlations of HD cell pairs during real arena light (RL) vs dark (RD, top) and RL vs VR-YawFree (bottom) from a representative session. **(B)** Heatmap of pairwise Kendall τ correlations of HD cell pairs from a representative session, RL vs RD (left) and RL vs VR-YawFree (right). **(C)** Boxplots showing the mean absolute τ correlations across RL, RD, and VR-YawFree sessions, each dot represents a single session. **(D)** Boxplots showing the absolute differences in correlations (Δ τ) RL – RD and RL – VR-YawFree. **(E)** Isomap projections of HD population activity for RL and VR-YawFree. The color represents heading directions. Persistent homology analysis was utilized to examine the topology of the HD manifold. The length of persistent Betti barcodes signifies the shape of the manifold, where (1,1,0) indicates the presence of the ring (right). (**F**) Off-ring manifold dynamics index quantification across sessions, comparing RL vs VR-YawFree. **(G)** Schematic of the custom head-fixation apparatus enabling mice to freely navigate the real-world arena while maintaining pitch and roll immobility (Hfix). **(H)** Pairwise correlations in RL, RL and RL-Hfix for an example session. **(I).** Mean pairwise correlations across RL, RD, and RL-Hfix sessions. **(J)** Δτ between RL-RD and RL-RL-Hfix conditions. (**K**) Isomap projections of HD population activity comparing RL and RL-Hfix conditions. (**L**) Same as in (**F**) but for RL vs RL-Hfix. *p < 0.05, **p < 0.01, ***p < 0.001.

We further explored the origins of these differences by examining population dynamics through dimensionality reduction and topological analysis techniques ^21^. To visualize the state space of the head direction network, we employed Isomap, a non-linear dimensionality reduction technique that preserves the intrinsic local structure of high-dimensional data^22^ (see^21^ for application of this technique to the study of the HD network). Input matrices in this analysis comprised the activity patterns of all simultaneously recorded neurons from each session, including both head direction cells and non-head direction cells. For visualization purposes, we projected the data onto the first two dimensions of the Isomap embedding, which revealed the topological structure of neural population activity. To quantify whether these embeddings formed a ring manifold structure, we conducted topological analysis using persistent homology, reporting Betti barcode numbers^21,23^. Despite the 2D visualization, this analysis was performed on five-dimensional Isomap embeddings to balance dimensionality reduction with preservation of topological features, allowing us to detect the presence and stability of ring-like structures in the neural activity space.

Topological analyses revealed that manifold dynamics deviated substantially from the ring structure in VR-YawFree, as evidenced by the absence of the H1 bar in VR-YawFree (**Fig. 2E**). This result manifested visually with the population activity shifting inside the ring rather than along its perimeter. To quantify the time spent “off-manifold,” we identified all samples within a recording session whose radius was below 30% of the 90th percentile radius for that session. Off-manifold indexes (0 = at manifold edge; 1 = at center) were consistently larger (indicating more off-ring manifold activity) in VR-YawFree (blue, **Fig. 2F**) compared to RL sessions (orange, **Fig. 2F**; p = 0.009).

To determine whether the observed alterations in correlations and manifold dynamics in VR-YawFree were attributable to the virtual reality environment or to head fixation itself, we developed a custom apparatus that immobilized the head relative to the body while still allowing the animal to freely locomote and turn during running (**Fig. 2G**). In this configuration, animals navigated the same real-world environment either with free head movement in all directions (RL), or with the head fixed to the body—eliminating head-on-body pitch, roll, and yaw rotations, while still providing normal vestibular yaw rotation signals through whole-body turning (RL-Hfix; a 3D-printed head-fixation apparatus mounted on wheels). Strikingly, we observed similar alterations in correlation structure and manifold dynamics in this condition, as we had observed in VR-YawFree (**Fig. 2H-L**). We observed weaker pairwise correlations than without head restraint (**Fig. 2H**, example sessions; **Fig. 2I**, Mann Whitney U comparisons to RL-Hfix, RL p = 0.008 and RD p = 0.01), and a strong remapping in RL-Hfix (**Fig 2J** abs (Δ1), p =0.0001). Ring manifold structure also collapsed in the RL-Hfix condition: the off-manifold index was consistently larger (more off-ring manifold activity) in sessions with the head-restrainer apparatus compared to regular arena sessions (**Fig. 2K and L**, p = 0.0002). Together, these results suggest that head fixation itself, and not the VR environment, alters HD population dynamics.

### Off-ring manifold dynamics emerge during stationary epochs

To further explore what drove off-ring manifold dynamics, data were segregated into stationary (*Stat*) and running (*Run*) epochs, with the latter being defined by a linear speed exceeding 5 cm/s (500ms bins; see Methods). This analysis revealed that locomotion significantly modulates ring manifold dynamics in the VR-YawFree environment, but not in RL. Both the length of Betti barcode H1 (a topological measure of ring structure) and manifold decoding error showed significant differences between *Stat* and *Run* epochs in VR-YawFree (**Fig. 3A**, example sessions; **Fig. 3B**, top: Wilcoxon-signed rank test in VR-YawFree: H1 barcode length, p = 0.002, n = 15 sessions; bottom: decoding error: p = 6.29e-5, n = 15 sessions. For RL, p = 0.45 and p = 0.23, respectively).

**Figure 3.**
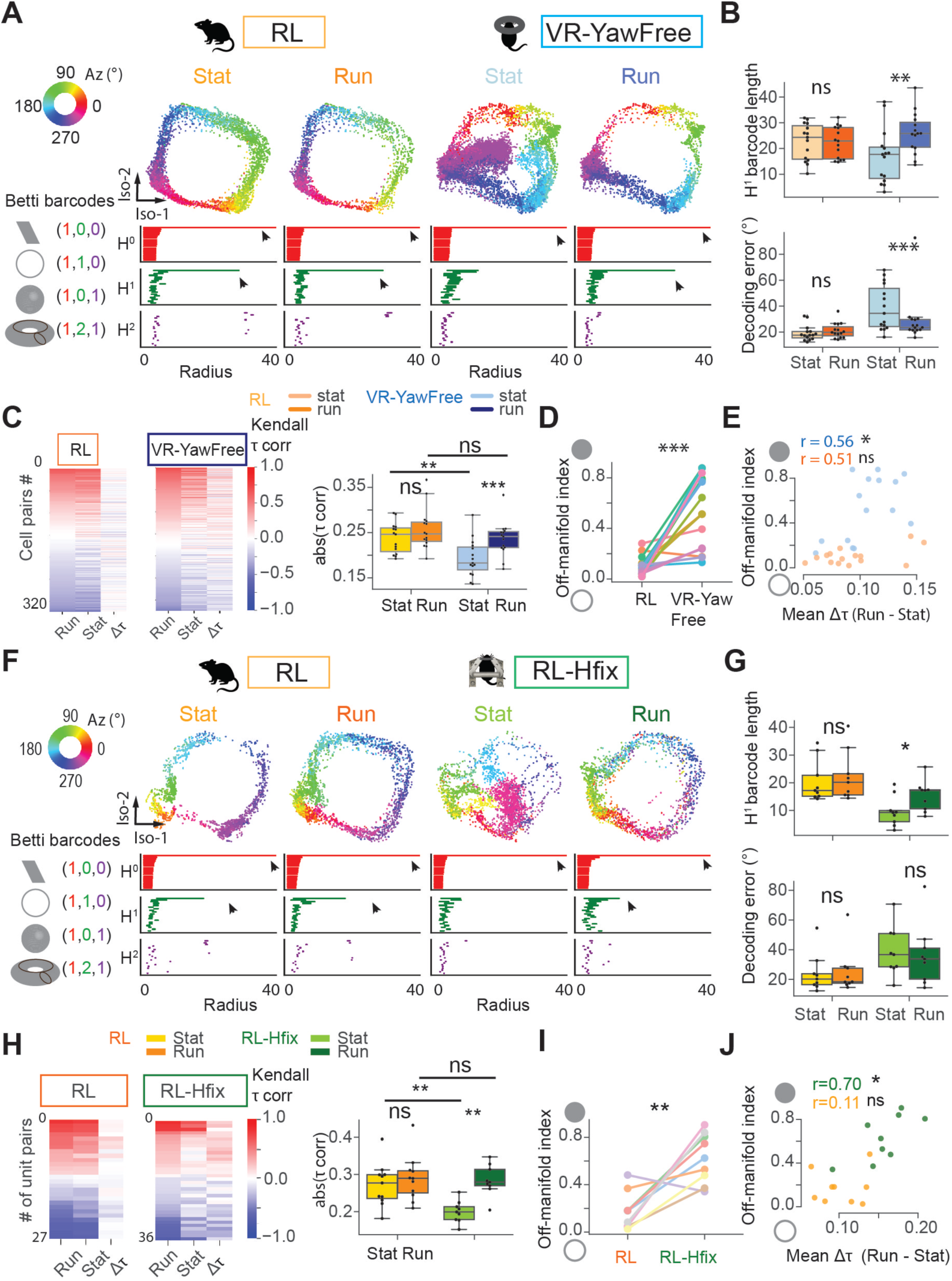
Off-ring manifold dynamics emerge during stationary epochs. **(A)** Isomap projections during *Stat* and *Run* epochs for a representative RL (left) and VR-YawFree (right) session. The color represents the animal’s head direction. Arrows indicate the presence of the persistent bar. (**B)** Boxplot of H^1^ barcode length (top) and Isomap decoding error (bottom) during *Stat* and *Run* epochs. **(C).** Left panel: Heatmap of pairwise Kendall τ correlations from a representative RL session (left) and during VR-YawFree(right) during Stat and Run epochs. Right panel: Boxplots show the mean absolute τ correlations during *Stat* and *Run* across environments, each dot represents a single session. **(D)** Off-manifold index for *RL* and *VR-YawFree* sessions during stationary periods: a higher value indicates increased off-manifold activity. **(E)** Scatter plot of off-manifold index vs. the mean absolute Δτ (Run - Stat), each dot represents a single session. **(F-J)** Comparison of freely moving mice with (RL-Hfix) and without head restraint (RL), showing similar disruption of ring dynamics during immobility in head-fixed condition. *p < 0.05, **p < 0.01, ***p < 0.001.

The same conclusions were reached when using an off-manifold index (0 = at manifold edge; 1 = at center; see Methods): off-ring manifold activity was stronger during stationarity (*Stat)* in *VR-YawFree* compared to *RL* conditions (**Fig. 3D**, p = 2.32e-5); and this was consistent across mice (**Figure S2**). Notably, mean firing rate of HD cells did not change, suggesting that observed differences are due to network dynamics (**Figure S3**). Thus, a locomotive state is required for normal HD population dynamics in head-fixed animals, even if single HD cell tuning appears intact.

Pairwise correlations between HD cells were also differentially modulated by locomotion across environments. While correlations remained relatively stable across *Stat* and *Run* in RL (**Fig. 3C**, left: p = 0.05), they were significantly reduced during immobility in *VR-YawFree* (**Fig. 3C**, right: p = 2.4e-04, n = 15 sessions). Direct comparison between conditions revealed significantly lower pairwise correlations during stationary periods in VR-YawFree compared to RL (**Fig. 3C**, p = 0.006), while correlations during running periods were comparable across RL and VR-YawFree (**Fig. 3C**; p = 0.21; p-values corrected using Benjamini/Hochberg method).

To investigate putative mechanisms underlying differences in head-fixed HD dynamics during (im)mobility, we examined whether the extent of network remapping (measured as the differences in pairwise correlations between RL and VR-YawFree) predicted the degree of off-manifold activity. For each recording session, we calculated the difference in Kendall correlations between *Stat* and *Run* epochs, and found this metric significantly correlated with the off-manifold index in VR (**Fig. 3E**, Pearson correlations p = 0.03). This relationship indicates that sessions exhibiting greater differences in correlation structure between behavioral states also displayed more pronounced off-manifold dynamics during stationary periods.

We next investigated whether similar on- and off-manifold dynamics are observed in mice exploring the real world. Indeed, off-ring manifold dynamics were observed during stationarity in RL-Hfix but not in RL, mirroring our observations in the VR-YawFree environment (see **Fig. 3F** for examples). This was particularly true when indexed via H1 barcode length (**Fig 3G**, top: RL, p = 0.13; RL-Hfix, p = 0.01), but also true when indexed via manifold decoding error (**Fig. 3G**, bottom: Δ decoding error *Stat* vs. *Run* in RL = 1.8 degs, and 3.0 degs (i.e., 1.66 times ΔRL) in RL-Hfix, p = 0.057, due to large variances). Quantification of off-manifold activity revealed significantly greater deviations from ring-like structure during immobility in real-world head-fixed periods compared to unrestricted periods (**Fig. 3I**, p = 0.004).

Similarly, pairwise correlations during stationary periods were significantly reduced in RL-Hfix compared to both their respective running epochs (**Fig. 3H**, rightmost, p = 0.007) and to stationary epochs in RL (**Fig. 3H**, right, green vs. yellow, p = 0.007). Moreover, as with the *VR-YawFree* condition, the degree of off-manifold population activity in *RL-Hfix* strongly correlated with the magnitude of changes in pairwise correlations between running and stationary states (**Fig. 3J**, RL: r = 0.11, p = 0.79; RL-Hfix: r = 0.70, p = 0.04). In contrast, HD single cell analysis showed that peak firing rate of HDC during stationary epochs was not significantly different between RL-Hfix and RL conditions, suggesting that observed differences are due to the network dynamics, and not unit activity (**Figure S4**). The selective increase in off-manifold frequency in stationary periods during head fixation (but not when the head was free to move) was consistent across animals (**Figure S5**). Collectively, these findings suggest that stationary and locomotion periods represent distinct population states in head-fixed animals, with stationary periods driving the disrupted network correlations and off-ring manifold dynamics observed during head fixation. The similarity between the effects observed in VR-YawFree and RL-Hfix conditions suggests that head fixation itself—rather than the virtual environment—is the primary driver of these altered dynamics. Most importantly, the results suggest that locomotion is able to rescue the impairments in HD network dynamics caused by head fixation.

### Linear velocity predictively rescues HD population dynamics during head fixation

Next, we investigated whether linear velocity, angular velocity, or both were necessary for rescuing ring-like population dynamics of HD cells during head-fixation. Remarkably, we observed that linear velocity alone – without the typically accompanying changes in angular velocity – was sufficient to restore on-ring manifold activity. Further, changes in manifold radius consistently preceded movement onset by approximately 550ms (**Fig. 4A** for an example, and **Fig. 4B** for averages). Thus, it is the prediction of linear velocity (and putatively it’s concomitant acceleration) onset that drives HD network stabilization onto the ring-like structure in head-fixed mice. A cross-correlation analysis between manifold radius and running speed confirmed this temporal relationship across transitions (n = 25), with the distribution of lags centered at approximately - 550ms (**Fig. 4B** bottom, i.e., radius increase leads motor onset). This temporal precedence strongly supports a predictive mechanism where motor efference, or anticipation of re-afference, of velocity or acceleration signals influence HD network dynamics well before the animal initiates movement.

**Figure 4:**
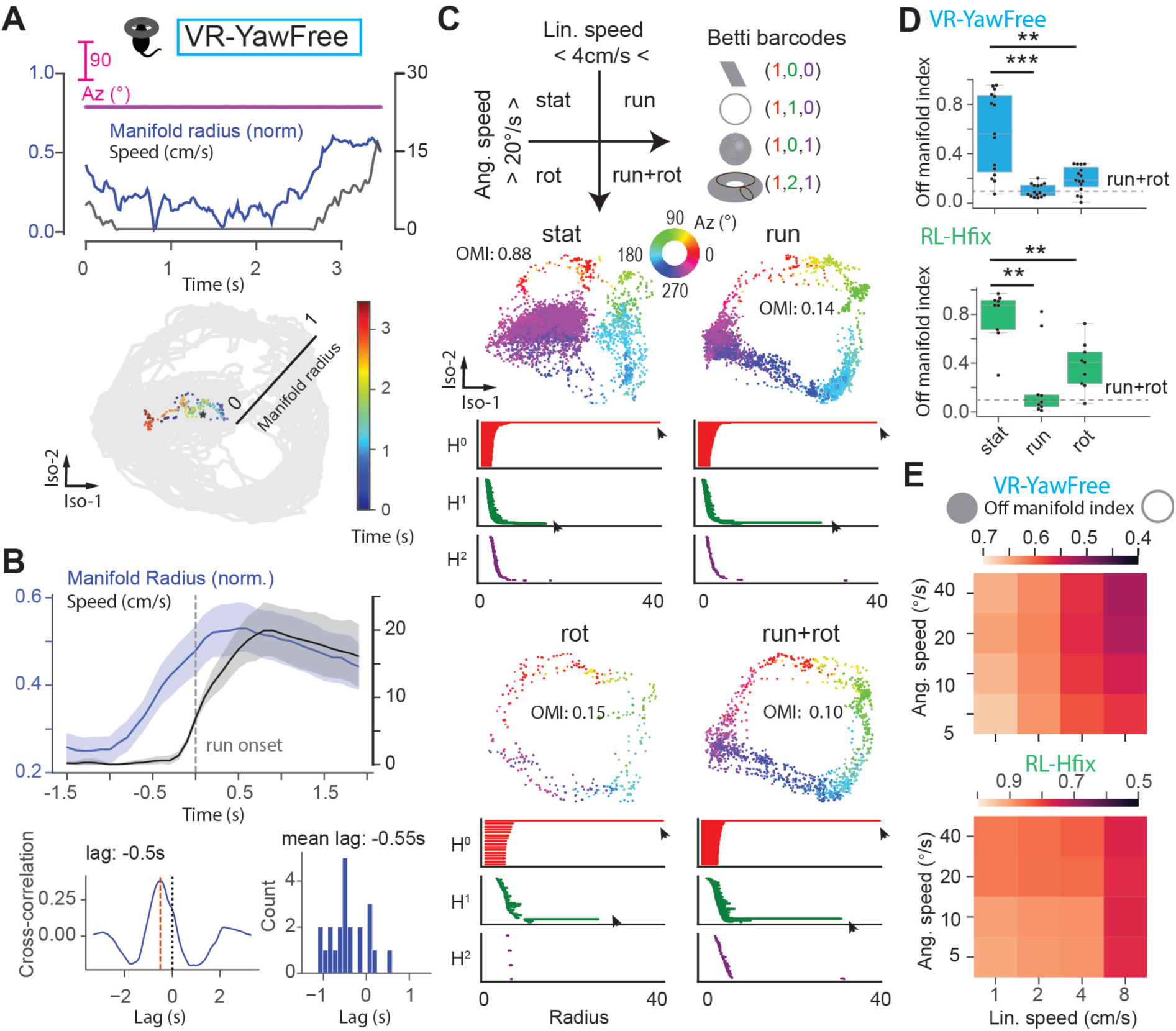
Prediction of linear velocity rescues HD population dynamics during pitch and roll fixation. **(A)** Representative example showing transition from stationary to running periods without head rotation (i.e., constant azimuth). The color gradient (bottom) shows the temporal progression, demonstrating how neural activity trajectories return to the ring manifold periphery as movement begins, despite the absence of head rotational cues. **(B)** Temporal relationship between linear speed and manifold dynamics. Top panel: Population-averaged traces of linear speed (black) and manifold radius (blue) aligned to running onset (n = 25 transitions, each of these without concomitant change in azimuth). Shaded regions represent ±1 SEM. Bottom left: Cross-correlation between speed and radius changes. Bottom right: Distribution of temporal lags calculated individually for each transition epoch. **(C)** Isomap projections along with Betti barcodes across different epochs. **(D)** Boxplots of the off-manifold index (larger the number = more off-ring activity) across different epochs for VR-YawFree (top) and RL-Hfix (bottom) environments. The dotted line represents the reference value of off-manifold index from the run+rot condition. **(E)** The off-manifold index was computed for various combinations of linear and angular velocity thresholds for VR-YawFree (top) and RL-Hfix (bottom). *p < 0.05, **p < 0.01, ***p < 0.001.

Next, we categorized animal behavior during VR-YawFree into four distinct epochs based on mean linear and angular velocity: stationary (*Stat*: linear velocity < 4 cm/s and angular velocity < 20°/s), running without rotation (*Run*: linear velocity > 4 cm/s and angular velocity < 20°/s), rotation without running (*Rot*: linear velocity < 4 cm/s and angular velocity > 20°/s), and combined running with rotation (*Run+Rot*: linear velocity > 4 cm/s and angular velocity > 20°/s; **Fig. 4C**). We found that off-ring manifold dynamics occurred primarily during stationary periods, and that either linear-or angular-velocity alone were sufficient to recover normal ring manifold structure (**Fig. 4D**; normalized by *Run+Rot* values, which are defined at y = 0.1). In fact, linear-only velocity periods result in off-manifold indices that were more dissimilar to the stationary period (p = 0.002) than the angular-only (i.e., rotation) periods were dissimilar to the stationary periods (p = 0.004; **Fig. 4D** top). We observed similar results in RL-Hfix, where either linear or angular speed alone was sufficient to maintain normal ring manifold structure (**Fig. 4D** bottom, *Run* p = 2.3e-5, *Rot* p = 0.003). Notably, during angular-only periods, heading direction changes, while during linear-only periods it remains constant. Despite this difference, both movement types—including linear velocity where heading is static—produce strong ring-like manifold structure in the HD network.

To ensure the robustness of our findings, we systematically varied the threshold values used to categorize periods as stationary, running, rotation, or running and rotating. This sensitivity analysis confirmed that our results remained qualitatively consistent across a range of threshold values: off-ring population activity predominantly occurred when mice were immobile, and that linear speed had a greater impact than angular velocity in determining on- vs. off-manifold dynamics (**Fig. 4E**; shades of red change predominantly along columns than rows). Together, these results suggest that HD population dynamics that are impaired during head-fixation are rescued to their on-ring manifold configuration prior to the onset of linear velocity; i.e., in a predictive manner.

### Reduced directional modulation of head-direction cells during off-ring manifold activity

To gain further insight into what is causing the off-ring manifold population activity, we focused on transitions from running toward stationary periods (*Stat*). We hypothesize that a reduction in HD cell firing rates at their preferred direction may contribute to off-ring manifold activity. To test this hypothesis, we categorized HD cells based on their preferred direction (within or outside the range of the current azimuth ±45°) and examined neural behavior as the population dynamics moved toward the center of the ring-like manifold (**Fig. 5A**).

**Figure 5:**
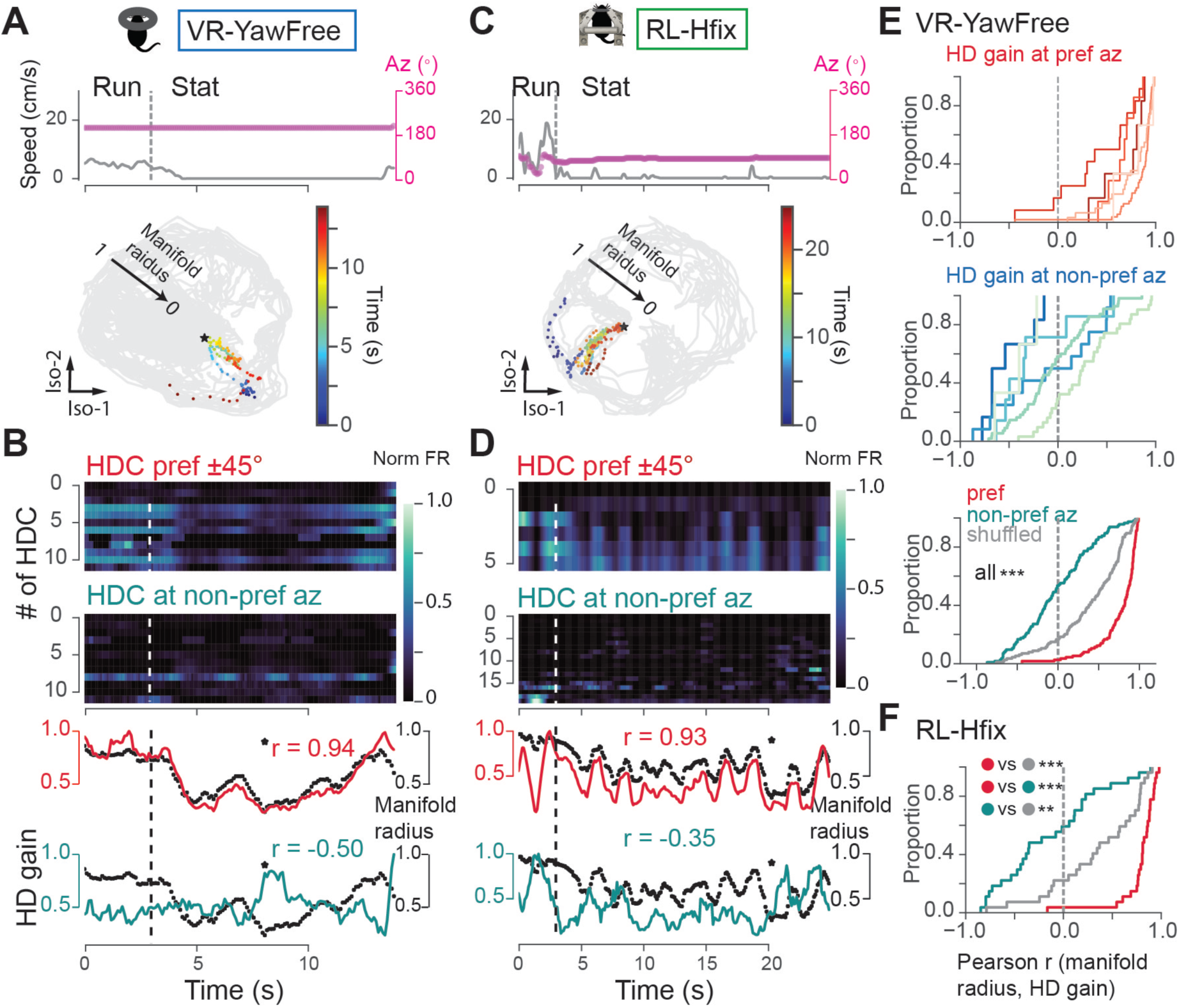
Reduced directional modulation of head-direction cells during off-manifold activity. **(A, C)** Representative transitions from running to stationary periods (no angular velocity) during VR-YawFree and RL-Hfix sessions. Top: Linear velocity (in gray) along with the animal’s heading (in magenta). Bottom: Manifold trajectory during transition from a running to a stationary epoch (color gradient signifies the progression of time). **(B, D)** Heatmaps display normalized firing rate of HD cells with a preferred direction within ±45° (top) and outside (bottom) of the animal’s current heading. Line plots depict mean population response (HD gain) for HD cells preferring the current azimuth (top) and those not preferring it (bottom), superimposed with the normalized manifold radius (black, 0 at the center, 1 at the edge of the ring manifold). **(E)** Cumulative distributions of Pearson’s r correlations between the normalized manifold radius and HD gain of HD cells preferring (top) and those not preferring (middle) the animal’s current heading. Each curve represents a different mouse. Data pooled across all animals are compared to a shuffled distribution (bottom). **(F)** Same as in **(E,** bottom) but for the RL-Hfix condition. *p < 0.05, **p < 0.01, ***p < 0.001.

The results show a decrease in HD cell firing at their preferred direction during stationary periods in VR-YawFree. To quantify the relationship between HD cell firing and manifold radius, we computed HD gain as the normalized mean firing rate of HD cells. We observed a positive correlation between manifold radius and HD gain at preferred directions, indicating an increase in HD gain as the manifold radius increased, and vice-versa. Furthermore, we observed a negative correlation between manifold radius and HD gain at non-preferred directions, suggesting reduced directional modulation of HD cells during off-ring manifold activity (**Fig 5B**). These findings were consistent across mice (**Fig. 5E**, r correlations between manifold radius and HD gain at preferred vs not-preferred direction, p = 1.69e-23, n = 220 stationary epochs).

Similar results were obtained for the RL-Hfix condition (**Fig. 5C-D**): During stationary periods, HD cell activity decreased at preferred directions, and increased at non-preferred directions (**Fig. 5F**, Pearson correlations between manifold radius and HD gain at preferred vs not-preferred direction, p = 3.24e-10). Collectively, these findings indicate that a reduced directional modulation of HD cell firing during stationary epochs (decreased at preferred headings and increased at non-preferred) results in off-ring manifold activity in both the VR-YawFree and RL-Hfix conditions.

### Perturbation of recurrent connectivity in HD ring attractor model recapitulates experimental effects of head-fixation

Since the degree of off-ring manifold dynamics in head-fixed mice was found to correlate with the magnitude of changes in pairwise HD cell correlations between *Run* and *Stat* epochs in both VR-YawFree (**Fig. 3E**) and RL-Hfix (**Fig. 3J**), via computational modeling we tested how changing the correlation structure would affect the manifold dynamics. We employed a previously published head direction continuous ring attractor model^24,25^. The model possesses the following key features: (1) visual landmark signals that activate HD cells anchoring their firing to the external environment; (2) HD cells are recurrently interconnected, amplifying the activity of neighboring cells to form a persistent bump of activity; (3) vestibular cues, primarily mediated by velocity cells, reinforce connections to steer the HD signal and enable the integration of angular velocity over time (**Fig 6A**).

**Figure 6.**
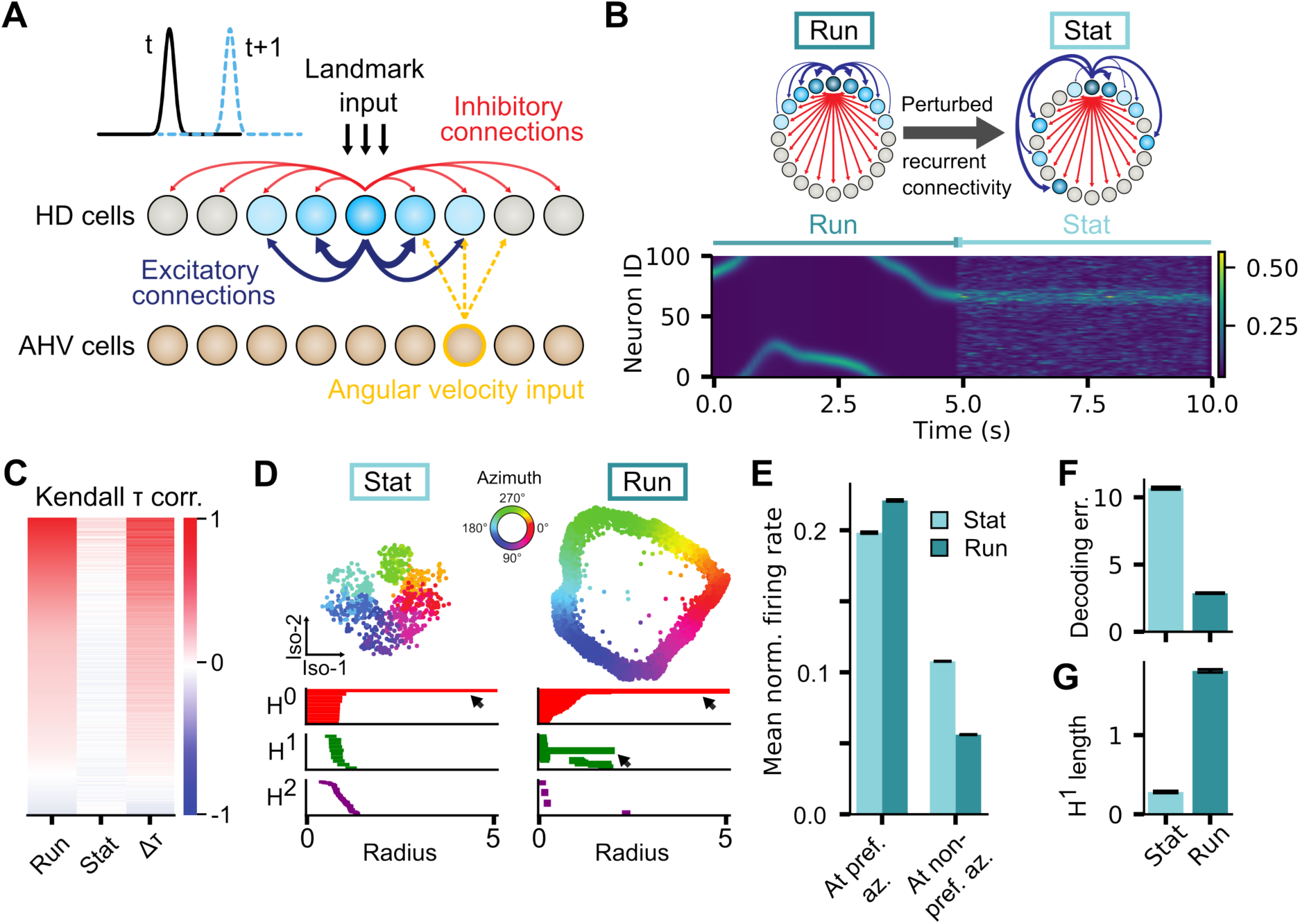
Perturbation of recurrent connectivity in HD ring attractor model recapitulates experimental effects of head-fixation. **(A)** Schematic of the ring attractor model. HD cells are activated by a visual landmark signal. Recurrent connections among neighboring cells amplify activity, forming a sustained “hill” of activity. Self-motion velocity signals, mediated by velocity cells, reinforce connections in a certain direction, enabling integration of velocity over time. **(B).** Top: Schematic of a random shuffling of the Gaussian recurrent connectivity profile, hypothesized to disrupt the pattern of connection strength between *Run* and *Stat* states. Bottom: Simulated HD cell responses. Note HD cell firing at non-preferred directions during Stat epochs. **(C)** Simulated Kendall tau correlation of the firing activities between pairs of HD cells for the two states. **(D)** Isomap projections during Stat (left) and Run (right) epochs, with corresponding Betti barcodes. Note that the ring topology is only present during Run, but not Stat periods. **(E)** Normalized mean firing rates of HD cells at preferred direction and outside that range during *Run* and *Stat* periods. **(F)** Decoding error from HD cell population activity during *Run* and *Stat* periods. **(G)** H^1^ Betti barcode length of the state space of HD population activity during *Run* and *Stat* periods. Data represented as mean ± SEM (n=5 simulations).

We simulated 100 HD cells, subjected to a random angular velocity trajectory to represent mouse head movements. This velocity trajectory was employed to activate the appropriate velocity cells, influencing the HD signal in the intended direction. The visual landmark input was based on actual azimuth, directly stimulating neurons with similar azimuth preferences. In our simulations, the HD cells within the network were recurrently interconnected, forming a Gaussian connectivity profile where neighboring cells (with similar preferred directions) had stronger connections compared to cells that were more distant. However, in stationary states characterized by the absence of head movement (*Stat*), we disrupted this connectivity pattern (**Fig. 6B**). This disruption led to an altered correlation structure during stationary periods, consistent with our experimental observations (**Fig. 6C**). Isomap visualization revealed changes in population activity, mirroring experimental data (**Figure 6D, D-G**). These changes resulted from reduced directional modulation of HD cells during stationary epochs, involving a decrease in activity among HD cells at their preferred azimuths, and a concurrent increase in activity among HD cells at non-preferred azimuths (**Fig. 6E**). We obtained similar results when decoding heading (**Fig. 6D**) or examining H1 Betti barcode lengths (**Fig. 6G**). These findings suggest that the pairwise correlation structure of the HD network is crucial for proper ring-manifold dynamics, and that it is this structure that is impaired during head immobilization, and which is recovered in anticipation of future movement / re-afference.

## Discussion

Our findings challenge a traditional view wherein the head-direction (HD) network is a passive, continuous integrator. Instead, the HD system appears to operate as an active, state-dependent estimator. During head fixation, immobility alters the HD ring attractor, whereas onset of locomotion predictively recovers its previous state. Critically, these changes were observed during both VR and real-world exploration, emphasizing the generality of the finding. Thus, ring-attractor models must be revised to include behavioral state and anticipatory motor cues accordingly.

Our restraining apparatus during exploration in the real-world immobilized head-on-body movements in all three-dimensions. However, in VR, mice could execute active yaw movements, and only pitch and roll were disabled. Under these conditions, the HD population covariance was severely disrupted and the manifold lost or altered its ring structure, even though individual cells remained tuned to direction (Fig S2 and S4). One explanation of these findings was that the missing tilt signals removed a crucial stabilizing reference. Indeed, HD cells are known to encode three-dimensional orientation with respect to gravity^8,9,26^. Similarly, animals with impaired otolithic (gravity) input show unstable HD firing^27^, and arguably even subtle head tilts (e.g. when rats are suspended in VR via a trunk harness as opposed to head-fixation^20^, help stabilize HD activity. In our data, the absence of pitch/roll cues in head-fixation likely deprived the network of this graviceptive anchor, promoting the “off-ring” activity we observed.

Crucially, we showed that it was head-fixation, and not the VR environment *per se*, that drove HD population-level changes. In a real-world setting where mice were head restrained (head-fixed on a wheeled apparatus), we saw the same disruption of pairwise correlations and apparent collapse of the ring manifold. Moreover, this collapse was highly state-dependent: it occurred during stationary periods. When head-fixed animals began to run (even on a treadmill), the ring dynamics were rapidly restored. In contrast, freely moving and freely head-tilting mice (in either lit or dark arenas) maintained stable ring manifolds whether they were still or running. Thus, the HD network appears to switch between two modes: “off-ring” mode during head-fixed immobility, and a normal ring mode during or before locomotion.

A key insight is that active locomotion alone can recover HD population dynamics. In our VR-YawFree recordings, increases in manifold radius actually preceded the onset of linear running speed by ∼550 milliseconds. This indicates that the network was responding to a predictive motor signal rather than sensory feedback or arousal from actual motion. In effect, an efference-copy of the upcoming locomotor command (or an internal prediction of the resulting acceleration) was sufficient to push the HD population back onto the ring manifold. Notably, this rescue required at least one intact vestibular channel: when we eliminated yaw input entirely (VR-YawFix), the predictive stabilization was lost (**Fig. S6**). In VR-YawFix, directional tuning was completely lost and peak firing rates were markedly reduced, indicating a collapse of HD network dynamics. This suggests that locomotion alone exerts only a modulatory influence on the HD system and cannot sustain coherent population activity in the absence of matching vestibular input. Alternatively, the severe sensory conflict in VR-YawFix may actively disrupt integration of motor and vestibular signals, preventing locomotion-related inputs from restoring manifold stability.

At the level of single cells, head-fixed immobility caused reduced directional tuning changes: cells fired less at their preferred directions and more at their non-preferred directions. This pattern flattens the population activity and shifts it “off” the ring (overall firing rates remained similar, but the covariance structure changed). To understand this, we adapted a continuous attractor model^24^ by weakening the recurrent connectivity among HD neurons during stationary head-fixation. This perturbation reproduced the empirical effects: the simulated network’s activity collapsed toward the center of the manifold, and tuning curves showed the same reduced directional modulation. While other mechanisms (e.g. neuromodulatory changes or global inhibition shifts) could also contribute, these modeling results confirm that context-dependent changes in network coupling can account for the experimental observations.

Together, these results have important implications for how we think about spatial coding. They suggest that the HD attractor network, while capable of self-sustaining activity through its intrinsic recurrent dynamics, may require predictive motor input for robust stabilization against drift and noise. Traditional ring-attractor models^5–7^ demonstrate that recurrent connectivity alone can maintain a stable activity bump in the absence of input. However, our data show that when motor efference signals are altered (as in head-fixation), the network becomes vulnerable to destabilization—the heading representation degrades not because the attractor dynamics fail, but because the system lacks the continuous error-correction signals normally provided by anticipated self-motion. In practice, the HD network appears to operate in an active inference mode, using forthcoming motor cues as a predictive stabilization mechanism that counters accumulated noise and keeps the bump precisely anchored. Modeling frameworks will therefore need to integrate these state-dependent dynamics, incorporating motor efference-copy signals not as drivers of the attractor *per se*, but as essential stabilizing inputs that maintain representational fidelity during naturalistic behavior.

Looking forward, it will be valuable to pinpoint the neural sources of these predictive signals and state effects. For example, recordings from motor or vestibular regions (or manipulations of neuromodulatory pathways) during the real-world and virtual navigation with and without head-fixation could reveal how the efference-copy stabilizes HD dynamics. It would also be interesting to test whether similar context-dependent manifold changes occur in other spatial circuits (such as hippocampal place cells or medial entorhinal cortical grid cells) under analogous head-fixed conditions^13,14^.

In summary, our findings reveal the HD system as an active, predictive estimator whose stability depends on both behavioral state and context. Constraining the head during immobility disrupts its internal compass, whereas locomotion or even just an internal drive to move reinstates a coherent ring attractor. This challenges the notion of a purely passive HD integrator and points to a more dynamic model in which self-motion predictions play a crucial role in maintaining a stable sense of direction.

## Acknowledgements

This work has been funded by NIH DC004260, Simons Collaboration for the Global Brain (SCGB) SFI-AN-NC-GB-Culmination-00002796. A.P. was supported by Nazarbayev University under Faculty-development competitive research grants program for 2025-2027 (Grant No. 040225FD4718). J.N. was additionally supported by NIH NINDS R00NS128075, the Simons Collaboration in Ecological Neuroscience (SFI-AN-NC-SCN-00007276-10 and SFI-AN-AR-Pilot-00008952), CTSI 1UM1TR004405, and a Sloan Research Fellowship.

## Author Contributions

D.A., J.A., J.N., and A.P. designed research; A.P. and J.A., performed experiments; H.Z. performed modeling; S.C.S, J.N., and A.P. provided analytical tools; A.P. and J.A. analyzed data; A.P., J.N., J.A., and D.A. wrote the paper with contributions from all coauthors; D.A. supervised research.

## Material & Methods

### Animals

We used a total of nine C57Bl6/J mice (3–6 months old). Six mice participated in virtual reality experiments and six in head-fixed freely moving experiments. Three animals contributed to both conditions on separate days. All animals were ran in the freely-moving conditions in order to identify HD cells. We housed the animals in a reversed 12/12-hour light/dark cycle to ensure recordings occurred during their naturally active phase. We conducted all procedures in accordance with US NIH guidelines and with approval from the New York University Animal Welfare Committee (UAWC).

### Surgeries

The surgical procedure consisted of two steps. First, a head-bar implantation which allowed for head-fixation and behavioral training. Second, a craniotomy allowing for electrode implantation. In the first step, we first deeply anesthetized each mouse with 3–5% isoflurane and secured it in a stereotactic frame (Kopf Instruments) via earbars. We covered the eyes with ointment (LubriFresh P.M Major) and applied 4% lidocaine topically. Then, a midline scalp incision was made, and we removed the periosteum with hydrogen peroxide (H₂O₂). After aligning Bregma and Lambda to level the skull, we placed a CNC-machined head-bar and fixed it with C&B-Metabond adhesive cement (Parkell). We reinforced the implant with Ortho-Jet acrylic (Lang Dental), forming a protective crown with an oval window over the skull landmarks. We sealed this window with UV-curable glue (Norland NOA 61) to protect the skull from potential infections. Animals were given 4 days of rest after this surgery before starting water restriction and behavioral training.

In a second step, after the mice completed training (see below), we anesthetized them again with isoflurane and placed them in the stereotactic frame (as above). We removed the UV glue and re-leveled the skull by re-measuring Bregma and Lambda. Then, a grounding skull screw was inserted over and touching the cerebellum. We performed a craniotomy at anterior-posterior (AP) –0.8 mm, mediolateral (ML) ±0.7 mm, dorsoventral (DV) 3.8 mm (targeting anterior thalamus). We carefully removed the dura mater, and inserted a four-shank Neuropixels 2.0 (NP2.0) probe (medial shank reference). Before lowering, we dipped the shanks in fluorescent DiI (Invitrogen Vybrant DiI) to label the electrode tracks for postmortem histology. The probe was attached to a series of 3D-printed enclosures, which were then cemented onto the animal (custom 3D housing available upon request to). Next, we applied bone wax over the craniotomy to stabilize the electrodes. A copper mesh at the back of the implant provided additionally shielding and protected the craniotomy and flex cables from debris. We also added a support to permanently connect the NP2.0 flex cable to the headstage, reducing strain on the probe. Animals were allowed 3 days of rest after this surgery before recording. To reduce the total number of animal, we recorded from both hemispheres in each mouse. We first implanted the left side. If the mouse remained healthy and performing well, we later repeated the surgery to implant the right side. We verified all electrode locations postmortem via histology (see below).

### Virtual Reality Apparatus

We built a 2D virtual reality setup using four LCD monitors (Scepter E20, 1600×900 pixels) arranged in a 45×45 cm square around a 15 cm radius air-supported Styrofoam ball. We used a custom head-plate holder to mount a slim bearing (KA030CP0) directly above the ball, allowing the mice to rotate freely in yaw while preventing head pitch and roll (see ^14^ for a similar approach). The head holder sat 8 cm above the monitors. We fixed the yaw rotation of the ball to couple the mouse’s rotation to the bearing, as in previous setups ^14,20^.

We ran unity-based VR software on a Dell Precision 5820 (Windows 10) with an Intel i9-10940X CPU and two NVIDIA Quadro RTX 5000 GPUs. We developed the VR environment in Unity3D (2019.3.15f1), assigning one virtual camera per monitor to create the scene. We tracked the Styrofoam ball’s translation in the XY plane with two orthogonally mounted ADNS-9800 laser motion sensors (Tindle), one in front and one to the side of the ball. These sensors were connected via Arduino to Unity. We calibrated each sensor by normalizing the virtual distance traveled to the actual distance rolled on the ball. We measured the mouse’s head direction using a circular touch sensor (rotary potentiometer, Tinkersphere) connected to Arduino, which sent the current azimuth to Unity. For reference, when a mouse physically turned on the ball it simply brought a different monitor into view; this physical turn did not directly rotate the virtual scene.

In a subset of sessions, we clamped the bearing so that mice had to use the ball to rotate the virtual environment. In this mode, rotation was triggered when a mouse ran at an oblique angle relative to the forward axis. Specifically, we calculated the trajectory angle from the ratio of forward and lateral velocity. If this angle fell within ±30° of the lateral axis, we treated the movement as an intentional turn. During these turns, we suppressed translational movement and converted locomotion into angular rotation. We set the virtual camera’s yaw velocity as a linear function of the lateral velocity (gain of 400°/s per unit lateral velocity). The sign of the lateral velocity determined left vs. right rotation. We implemented a small dead zone to avoid unintended rotations and reset translational velocity to zero when a turn engaged. This scheme let mice initiate virtual turns by running at an angle on the ball, while forward locomotion (|forward| >> |lateral|) continued to produce forward movement through the virtual space.

We delivered rewards through a custom lick tube attached to the head holder and positioned within licking distance. An Arduino-controlled solenoid released the 10% sucrose solution by gravity. We calibrated the system by measuring dispensed volume versus valve opening time, ensuring that each trial delivered 5 µL of solution.

### Behavioral training

Mice were handled for 20 minutes daily starting one week before the first surgery. After 4 days of recovery from surgery, we began VR behavioral training. We started water restriction 3 days post-surgery to motivate the mice.

For the first two days of VR habituation, each mouse sat on the Styrofoam ball in darkness, and we delivered a 10% sucrose reward every 20 seconds. After the mice were familiarized with the setup, we turned on the airflow, which suspended the Styrofoam ball, and screens. The mice then saw a moving blue beacon, and when it intercepted the beacon, we gave it sucrose paired with a tone. The ball’s movement was initially locked to one dimension (forwards/backwards). In this phase, the mice earned a reward by running past a target, beaconed location. Once a mouse achieved >60% correct trials, we gradually increased target distances up to 0.75 m. On average, mice learned to run on the ball within 5 days.

In the final training phase, we removed the 1D constraint so that the mice navigated a 2D open field. We continued rewarding them with 10% sucrose. We ensured each mouse consumed at least 1 mL of liquid per day by supplementing with water if needed (mainly during the first days of training until they earned ∼1 mL from rewards). The mice were always provided any supplement water at the end of the day to avoid linking thirst to poor performance. Overall, the full training protocol took about 6–8 weeks.

### Electrophysiological recordings

We recorded extracellular signals with four-shank Neuropixels 2.0 probes using SpikeGLX software (v.20201024). We used two different channel configurations: either the bottom 384 channels or the 384 channels above these. Configurations were chosen based on the number of head-direction cells identified during initial screening in a real-world open arena.

#### Open-field arena

We recorded neural activity in a custom black square arena (50×50 cm). Each session started with an 8-minute light period followed by 8 minutes of darkness. We placed a white cue card on one wall during the light session, to act as a landmark. Food crumbles were scatterned randomly in the arena before each session to encourage foraging. The arena was cleaned between sessions to remove scent cues.

We tracked the mouse’s position at 50 Hz with a Chameleon USB3 camera (FLIR, CM3-U3-13Y3M-CS, 640×512 resolution). We extracted the animal’s heading and position offline using DeepLabCut (v2.2; Nath et al. 2019). The particular model of DLC was trained with 150 frames with varied lighting conditions (using an NVIDIA Quadro RTX 5000 GPU). The model tracked four body parts: left ear, right ear, head/implant, and tail base. We computed head direction as the arctangent of the ear positions. We calculated linear speed from the animal’s position (mean of the two ears and tail base) normalized to the arena’s dimensions. We synchronized the video and electrophysiology by generating random TTL pulses with an Arduino and sending them to both the camera and our acquisition system. These two streams of pulses were then cross-correlated to find the delay (i.e., intercept) that resulted in the best match (r = 1). Further, drifts in time clocks were corrected given the slope of these pulse correlations. We acquired images and metadata with Bonsai software.

#### Virtual reality with free yaw rotations (VR-YawFree)

After completing the open-field recordings, we moved the mice to the VR setup for an active foraging task. The mice navigated toward blue beacons to obtain 10% sucrose rewards. Each session lasted 20 minutes. We ran two types of virtual environments. In the first, an “infinite world,” the mice could move freely with no boundaries. In the second, virtual walls confined navigation to a 50×50 cm area. For analysis, we used only the walled sessions, though we observed qualitatively similar behavior in the infinite world.

#### Virtual reality with full head-fixation (VR-YawFix)

In five additional sessions, we recorded head-direction dynamics in a 2D VR environment with the mice fully head-fixed. In this condition, the virtual view rotated directly with the ball’s movements. The mice performed the same target-pursuit task for 20 minutes per session. The final dataset included two sessions in the walled VR environment and three in the infinite world. As in VR-YawFree, head-direction dynamics were similar in both environments.

#### Head-fixation in an open-field arena

Separately from the VR experiments, we ran open-field sessions with head fixation. We designed a custom 3D-printed apparatus to restrict the mouse’s head movements while allowing it to move. The custom piece (available upon request) employed an identical head-fixation mechanism as in our VR, but was mounted on wheels allowing animals to yaw-rotate and translate across the arena. The mice acclimated to this setup for at least seven days before recording. During recording, each mouse explored the arena. We first ran an 8-minute head-free light session followed by an 8 minutes head-free session in darkness. Then repeated another 8-minute light dark sessions with the head-fix component attached. We affixed a red LED and an IR LED to the headstage to track the mouse’s position and head orientation. We captured and processed the video data with a Plexon recording system.

### Spike detection and sorting

We sorted spikes in MATLAB using Kilosort2.5 ^28^. After sorting, we manually curated the clusters with the Phy2 graphical interface. We excluded any unit that was classified as noise, had a mean firing rate below 0.5 Hz, showed abnormal waveforms, or appeared only during a single part of the recordings (e.g., only in VR or only in the arena).

### Histology

After the recordings, we perfused each mouse transcardially with 4% paraformaldehyde (PFA) and post-fixed the brain in 4% PFA for 24 hours. We then transferred the brain to 30% sucrose for 48 hours. We cut 40 µm coronal sections using a cryostat. We mounted the sections on glass slides and straightened them in phosphate-buffered saline (PBS). After drying, we cover-slipped the slides with Fluoroshield mounting medium containing DAPI (Abcam). The next day, we imaged the DiI-labeled electrode tracks with a Leica fluorescence microscope. We exported the electrode coordinates as CSV files, and alignment histology using SHARP-Track software ^29^.

### Analyses

#### Identification of HD cells

To define HD cells, we employed a permutation test, constructing tuning curves by averaging the firing rates of units within 20-degree bins and normalizing them according to the animal’s occupancy. In the process, we conducted 1000 permutations for each neuron, circularly shuffling the azimuth to generate a distribution of Rayleigh vector lengths. For a neuron to be classified as an HD cell, it was required to possess a Rayleigh vector length greater than 0.3, surpassing the 95th percentile of the null distribution of vector lengths.

#### Criteria for Population-Level Analyses

For population-level analyses, we required at least 10 simultaneously recorded head-direction cells (HDCs). Because recordings were obtained from four shanks of a Neuropixels probe, we applied the following inclusion criteria: (i) sessions were retained if at least 10 HDC clusters were detected on any single shank, and (ii) adjacent shanks were also included if they contained more than 4 HDC clusters during freely moving, lights-on (RL) sessions in the real arena. Based on these criteria, 15 sessions from 7 implantations across 6 mice were selected for comparing HD dynamics between real and virtual environments. Histological verification confirmed that in these implantations, 7/7 were located in the anterior thalamus (ATN), with 6/7 specifically in the anterodorsal nucleus (ADN). For the freely moving, head-fixed condition, 9 sessions from 6 implantations across 6 mice were included. Histology confirmed anterior thalamic placements for all 6 implantations, with 3/6 localized to the ADN.

#### Pairwise correlations

To assess coordinated firing between neurons, we calculated Kendall’s tau (τ) correlations for all cell pairs using binned neural activity with a 20-ms resolution. Kendall’s τ quantifies rank correspondence between two variables: values approaching +1 indicate strong co-firing (both cells active together), whereas values near –1 reflect strong anti-correlated firing ^30^. By comparing τ distributions across contexts (freely moving vs. VR) and behavioral states (stationary vs. running), we characterized the intrinsic structure of the HD network.

#### Persistent homology

To visualize the population dynamics of head-direction (HD) cells, we constructed two-dimensional (2D) embeddings of the high-dimensional population activity. Spike trains of HD cells were convolved with Gaussian kernel and downsampled to 10 Hz and square-root transformed to stabilize variance. The firing rate matrix was then projected into a lower-dimensional space using the Isomap algorithm (scikit-learn, 50 neighbors) ^31^, yielding 2D embeddings for visualization and decoding, and 5D embeddings for topological analyses. The manifold angle was computed as the arctangent of the two coordinates in the 2D embedding space. Persistent homology was performed on the first five embedding dimensions. Betti barcodes (0, 1, and 2) were extracted using the Ripser package ^32^. For embeddings with more than 100 features, we restricted analyses to features whose lengths exceeded the 95th percentile. The length of the Betti-1 barcode was used as a quantitative measure of the ring structure within the manifold, enabling comparisons across experimental conditions.

#### Decoding head direction from the manifold embedding

Head direction was decoded directly from the two-dimensional Isomap space by calculating the arctangent of the ratio between the two embedding coordinates, yielding a neurocentric angular estimate. To assess accuracy, we computed the decoding error as the average absolute difference over time between the animal’s true head angle and the angle derived from the manifold.

#### Off-manifold index (OMI)

To compare stationary versus locomotor states, we defined the off-manifold index as the proportion of stationary time points whose radius from the manifold centroid fell below the 10th percentile of the locomotion radius distribution. For comparisons across experimental conditions, we instead used a normalized criterion: the off-manifold index was defined as the fraction of points lying within 30% of the 90th percentile of the whole session radius distribution. This empirically chosen threshold reliably captured whether the interior of the ring manifold was occupied by neural activity or remained sparsely populated.

#### Cross-correlation analysis of manifold radius and linear speed

To examine the temporal relationship between neural manifold geometry and animal linear velocity, we computed cross-correlations between the radius of the Isomap embedding and the mouse’s linear speed during virtual reality sessions. Both signals were mean-centered, and their cross-correlation function was calculated at 10 Hz resolution. We selected transitions where: 1) mean linear velocity was less than 2 cm/s in the 0.5s window before movement onset; 2) mean velocity exceeded 5 cm/s in the 0.5s window after movement onset; 3) head direction remained stable (circular standard deviation of azimuth < 2°) for at least 1.5s before onset to eliminate confounding effects of angular velocity; 4) normalized manifold radius dropped below 0.3 during the identified epoch, confirming off-ring dynamics during immobility; and 5) the cross-correlation between manifold radius and linear velocity was statistically significant (p < 0.05; permutation test). The peak cross-correlation and its corresponding lag were extracted for each run epoch. To evaluate statistical significance, we implemented a permutation test in which the manifold radius time series was randomly shuffled 500 times to generate a null distribution of peak cross-correlation values. An observed peak was considered significant if it exceeded the 95th percentile of the null distribution (p < 0.05). For significant epochs, the lag associated with the peak correlation was recorded, providing an estimate of the temporal offset between changes in neural manifold radius and locomotor speed.

#### Quantifying off-manifold dynamics across linear and angular velocities

Stationary and running epochs were identified using a 0.5 s moving window and a linear speed threshold of 4 cm/s, while rotational epochs were identified with the same window and an angular velocity threshold of 20°/s. These indices were combined to define four behavioral states:

stationary without rotation (stat), stationary with rotation (rot), running without rotation (run), and running with rotation (run+rot). For each recording, the reference off-manifold threshold was defined as the 10th percentile of the radius distribution during the run+rot state, since this condition typically exhibited the most stable ring structure. The off-manifold index for a given state (e.g., stat) was then calculated as the fraction of time points in that state with a radius smaller than the run+rot threshold. This allowed direct comparison of manifold occupancy across behavioral states relative to a common baseline. To confirm robustness of state definitions, the analysis was also repeated across a grid of thresholds (linear speed: 1, 2, 4, and 8 cm/s; angular velocity: 5, 10, 20, and 40°/s), although results are reported using the default thresholds of 4 cm/s and 20°/s.

#### Analysis of head-direction cell dynamics during stationary off-manifold epochs

To examine the relationship between HDC activity and the structure of the neural manifold during immobility, we identified stationary epochs (linear speed < 5 cm/s, minimum duration ≥ 3 s) from each recording. For each epoch, the manifold radius was computed from the Isomap embedding of population activity and normalized to the maximum radius observed within the session. Epochs were considered “off-manifold” if the normalized radius fell below 0.5 for at least part of the stationary period. For each off-manifold epoch, we calculated the circular mean and variance of the animal’s head angle. Epochs with a circular standard deviation > 1 radian were excluded to ensure stable head orientation. HDCs recorded in the session were then split into two groups: (i) preferred-direction cells, whose tuning curves aligned within ±45° of the animal’s mean head direction during the epoch, and (ii) non-preferred cells, with tuning offsets > 45°. For each group, we normalized firing rates by the maximum activity of each unit and computed the population gain as the mean normalized firing rate across units. We then quantified the relationship between neural activity and manifold geometry by calculating Pearson correlations between the normalized manifold radius and the population activity of preferred and non-preferred HDCs. To establish a baseline, correlations were also computed for randomly selected subsets of HDCs (20 random draws per epoch). This procedure yielded correlation values (r) for preferred, non-preferred, and random HDC populations during stationary off-manifold periods, allowing us to assess how the stability of the ring manifold relates to the alignment of HDC firing with the animal’s orientation.

#### Ring attractor model

The HD model used is a recurrent attractor network with global inhibition. A group of neurons are interconnected through synaptic weights determined by a Gaussian function which relies on the distance between the agent’s states in the physical world (i.e. head directions) represented by these neurons ^24,25^. Neurons representing similar head directions exhibit stronger connections. The network adjusts its firing rates using the following dynamical equations based on the ’leaky-integrator’ model. The activation *h^HD^_i_* of each head direction cell 𝑖 evolves through the following equation:

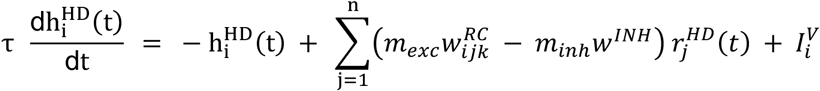

where r^HD^_j_ is the firing rate of head direction cell 𝑗 and *w^RC^_ijk_* symbolises the positive excitatory synaptic weight from head direction cell 𝑗 to cell 𝑖 modulated by the activated velocity cell 𝑘. Additionally, 𝑤^*INH*^ represents the global inhibition constant, indicating the impact of inhibitory interneurons. τ serves as the time constant regulating the system, while 𝑚_*exc*_ and 𝑚*_inh_* are scalar constants controlling the balance between excitation and inhibition in the system. 𝐼*^V^_i_* represents the visual landmark input received by head direction cell 𝑖 and it is characterized by a Gaussian response profile centered at the preferred head direction cell. The firing rate 𝑟*^HD^_i_* of cell 𝑖 is determined by passing the activation ℎ*^HD^_i_* through a sigmoid function:

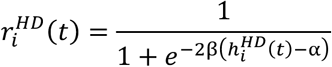

where α denotes the sigmoid threshold and β represents the slope of the sigmoid function. The standard parameters used for the simulations conducted in this paper are provided in Table 1.

**Table 1.** Model parameters. 𝑆𝑇𝐴𝑇 and 𝑅𝑈𝑁 refer to the different states during the simulation.

|  |  |
| --- | --- |
| $\tau$ | 1.0 s |
| $\alpha$ | 30 |
| $\beta$ | 0.022 |
| $m_{exc}^{RUN}$ | 4.68 |
| $m_{inh}^{RUN}$ | 3.12 |
| $m_{exc}^{STAT}$ | 13.52 |
| $m_{inh}^{STAT}$ | 3.12 |

In the simulations, the network consisted of 100 fully connected HD cells, each cell exhibiting a preference for a specific azimuth uniformly distributed from 0° to 360°. There were a total of 101 velocity cells: one cell represented no head movement, while the remaining 100 cells were evenly split to represent both positive and negative angular velocities. These velocity cells became active when the simulated agent experienced corresponding angular velocities. Subsequently, these active cells would adjust the recurrent connectivity between the HD cells in the simulation to propagate the hill of activity in a certain direction. During the stationary state, the pairwise correlation of firing activities among HD cells is perturbed by randomly shuffling *w^RC^_ijk_* and increasing 𝑚_*exc*_.

The decoding error of the model was computed as the absolute value of the difference between the real HD and the decoded HD. The decoded HD was calculated as a weighted average of the preferred directions of HD cells based on their firing rates at a specific time:

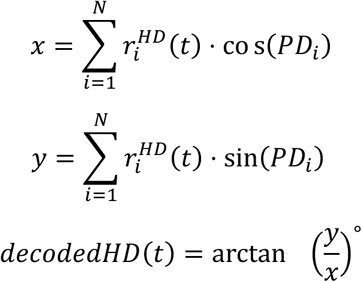

where 𝑟*^HD^_i_*(𝑡) is the firing rate of cell 𝑖 at time 𝑡, and 𝑃𝐷_*i*_ is the preferred direction of cell 𝑖 in radians.

### Statistics

The Mann–Whitney U test was used to compare two independent conditions (e.g., freely moving vs. VR). The Wilcoxon signed-rank test was applied to paired samples (e.g., stationary vs. running). To assess differences between two distributions, the two-sample Kolmogorov–Smirnov test was used.

## Supplementary figures

**Figure S1:**
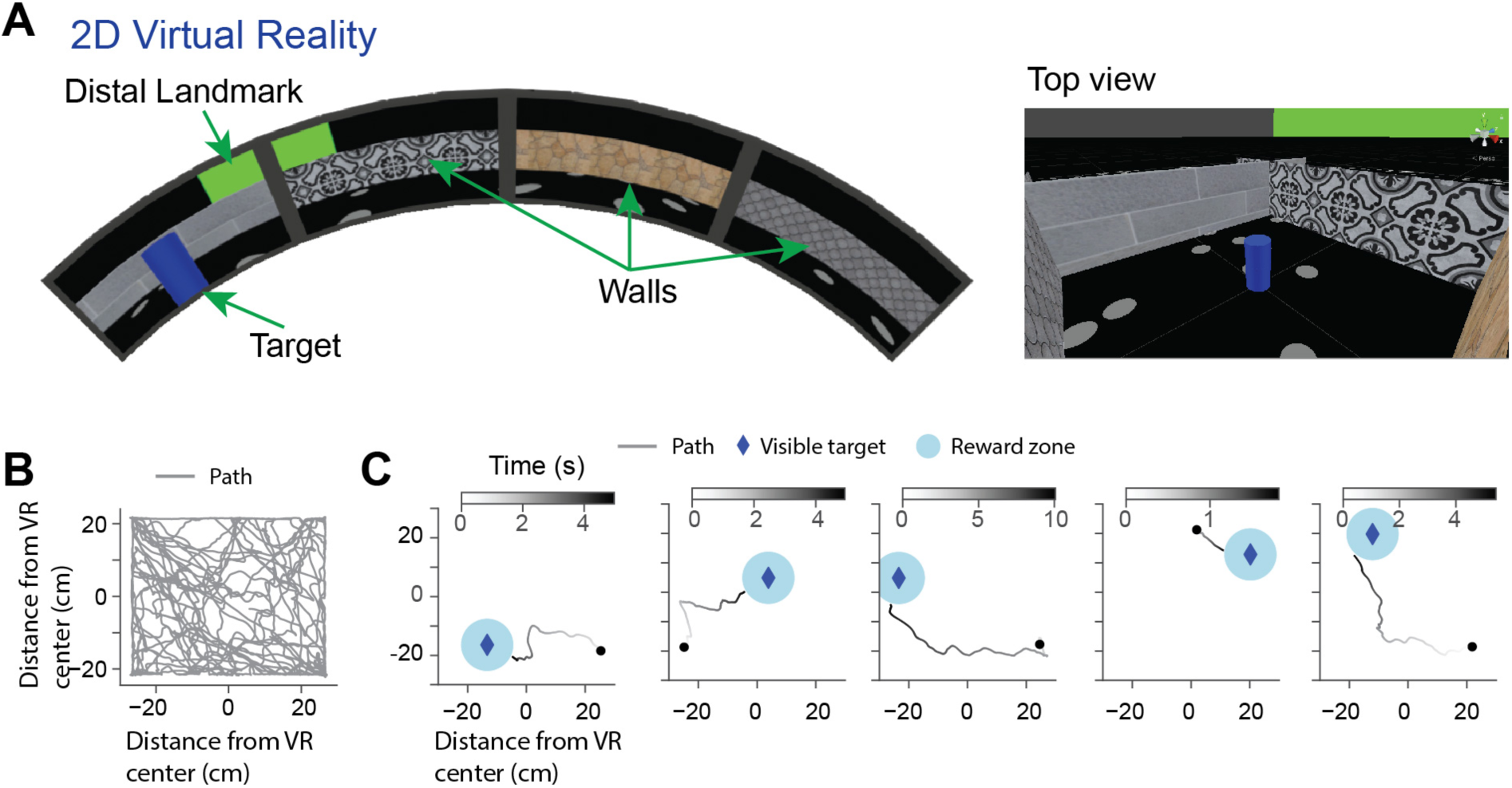
Target pursuing task in 2D VR. **(A, left)** An image of a 2D virtual square arena utilized in the task. Four monitors were arranged in a square configuration, providing complete azimuth coverage. **(A, right)** Top view of the virtual arena. **(B)** Representative example of mouse trajectory in a 2D VR. **(C)** Mice were trained to navigate to a blue beacon for a sugar reward. Five sample trials depict the trajectories of the animals as they navigate towards the target.

**Figure S2:**
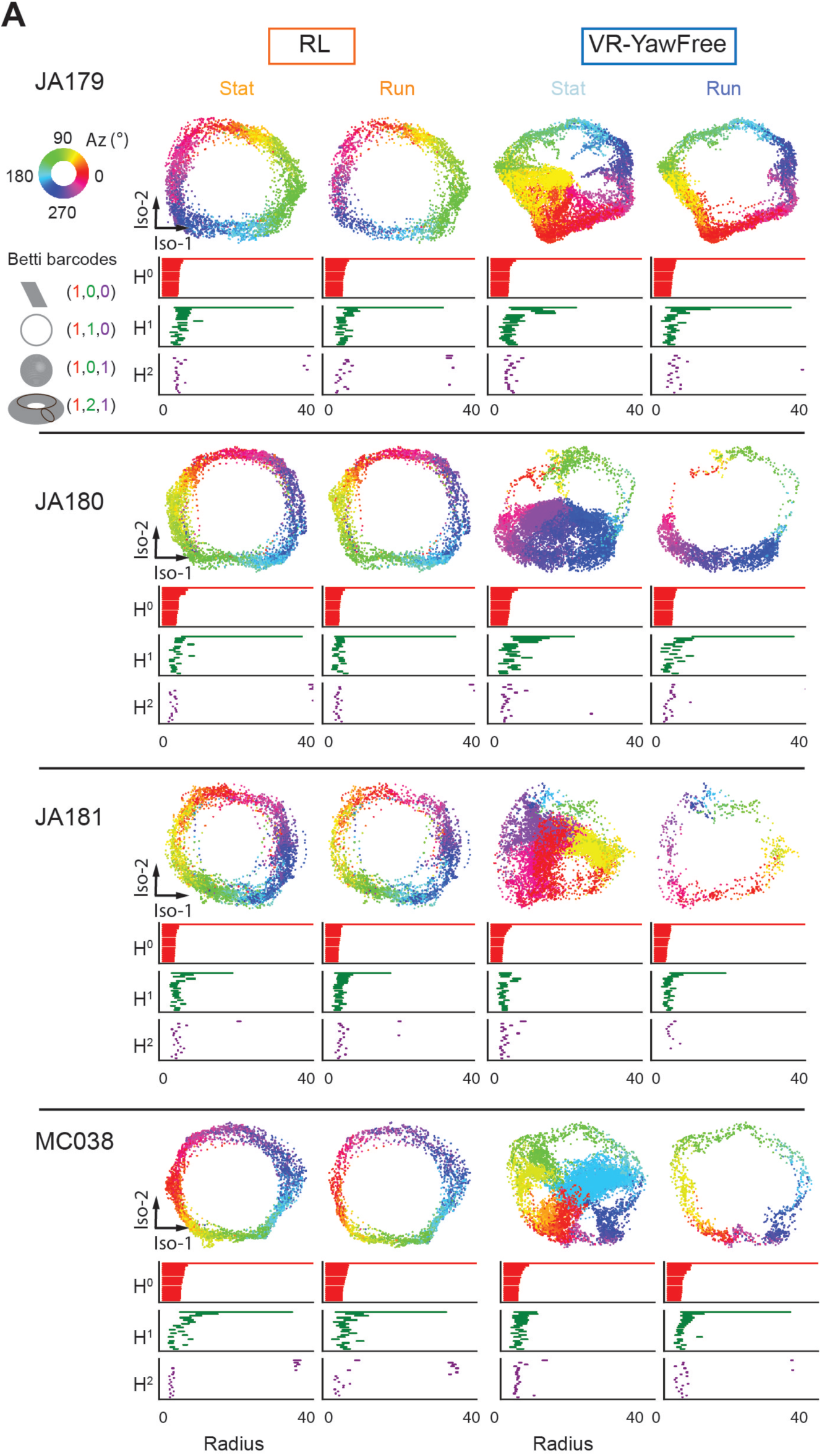
Off-ring dynamics in VR-YawFree across mice. **(A)** Example Isomap projections and corresponding Betti barcodes during Stat and Run epochs for Arena and VR-YawFree sessions. Each row represents a different mouse.

**Figure S3:**
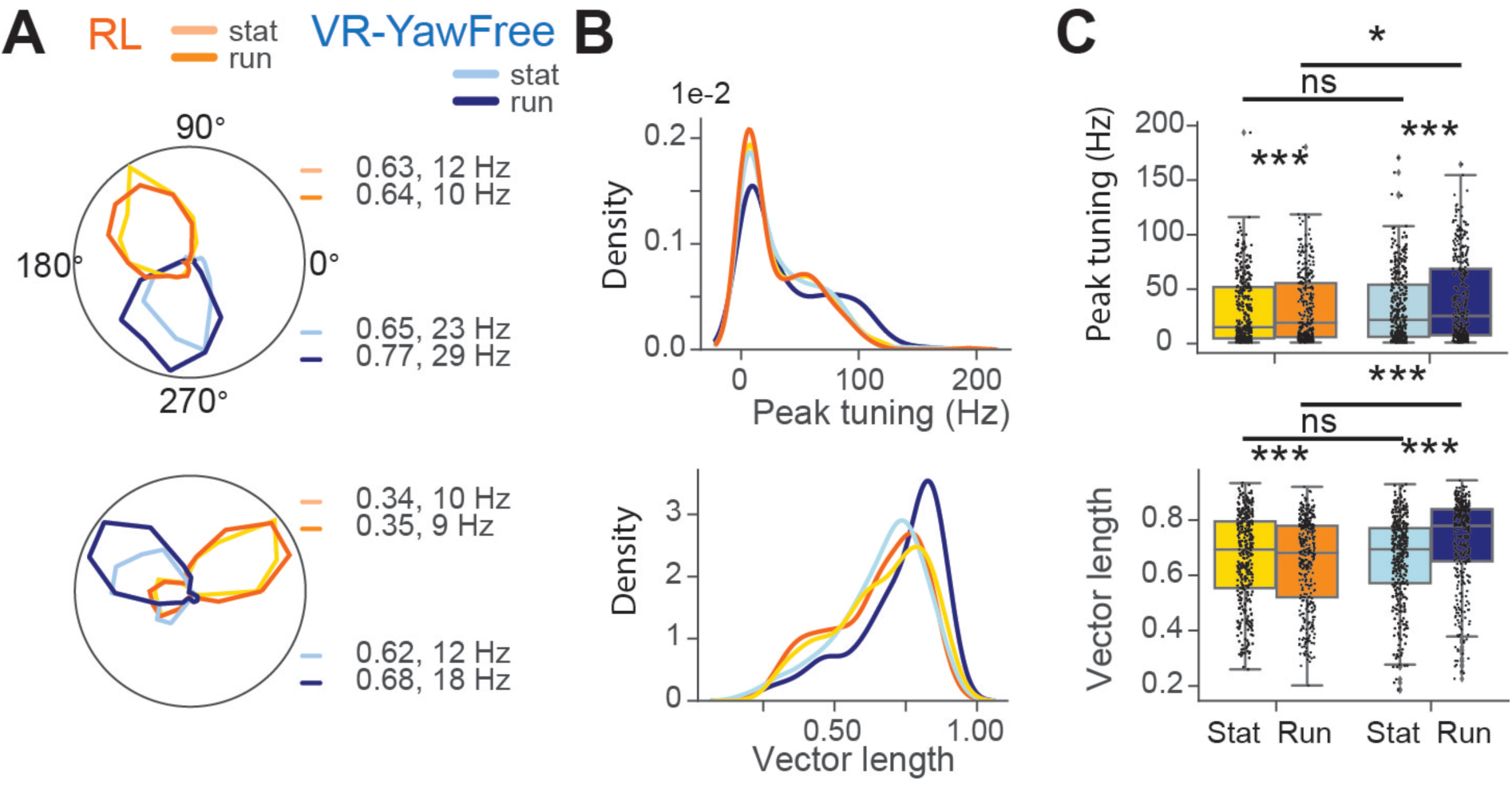
HD single cell properties are comparable between VR-YawFree and RL during stationary epochs. **(A)** Example HD cell tuning curves constructed from running (Run) and stationary (Stat) epochs for RL (orange) and VR-YawFree (blue). **(B)** Distribution of peak tuning curve amplitude (top) and Rayleigh vector length (bottom) across conditions. **(C)** Boxplot of tuning curve peak firing rates (top) and Rayleigh vector length (bottom). Wilcoxon-signed rank test peak tuning curve: Stat vs Run RL: median = 15.13 vs 19.05, W = 15957, P = 1.27e-15, n = 355 units; VR-YawFree: median = 21.74 vs 25.44, W = 10749, P = 1.18e-28, n = 365 units; Rayleigh vector length Stat vs Run RL: median = 0.69 vs 0.68, W = 22234, P = 1.74e-6, n = 355 units; VR-YawFree: median = 0.69 vs 0.78, W = 9049, P = 1.51e-33, n = 365 units. Mann Whitney U test peak tuning curve: Stat RL vs VR-YawFree, median = 15.13 vs 21.74, U = 65962, P = 0.22; Run RL vs VR-YawFree, median = 19.05 vs 25.44, U = 56747, P = 0.01, n = 355 vs 365 units; Rayleigh vector length: Stat Arena vs VR, median = 0.69 vs 0.69, U = 65962, P = 0.67; Run RL vs VR-YawFree, median = 0.68 vs 0.77, U = 46117, P = 3.30e-13, n = 355 vs 365 units, p values were adjusted using Benjamini/Hochberg method. *p < 0.05, **p < 0.01, ***p < 0.001.

**Figure S4:**
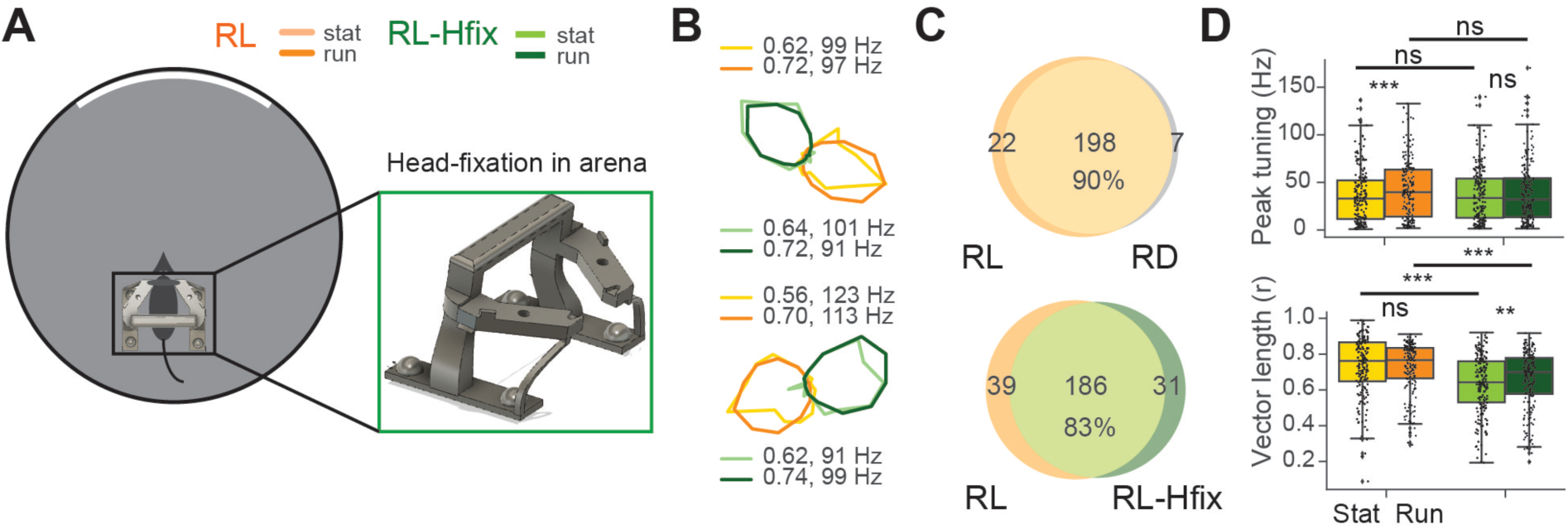
Running improves HD single cell tuning in RL with head restraint but not without it. **(B)** Example HD cell tuning curves constructed from Run and Stat epochs. **(C)** Pie chart of HD cells for RL vs RD (top) and RL vs RL-Hfix light (bottom). **(D)** Boxplot of tuning curve peak firing rates (top) and Rayleigh vector length (bottom). Wilcoxon-signed rank test peak tuning curve: Stat vs Run RL: median = 32.98 vs 39.66, W = 3810, P = 7.37e-11, n = 187 units; RL-Hfix: median = 33.60 vs 31.82, W = 8274, P =0.13, n = 197 units; Rayleigh vector length Stat vs Run RL: median = 0.76 vs 0.77, W = 7568, P = 0.10, n = 187 units; RL-Hfix: median = 0.64 vs 0.70, W = 7141, P = 0.001, n = 197 units). Mann Whitney U test peak tuning curve: Stat RL vs RL-Hfix, U = 18187, P = 0.13; Run RL vs RL-Hfix, U = 19942, P = 0.83, n = 187 vs 197 units; Rayleigh vector length: Stat RL vs RL-Hfix, U = 25182, P = 1.99e-9; Run RL vs RL-Hfix, U = 23405, P = 9.06e-6, n = 187 vs 197 units, p values were adjusted using Benjamini/Hochberg method. *p < 0.05, **p < 0.01, ***p < 0.001.

**Figure S5:**
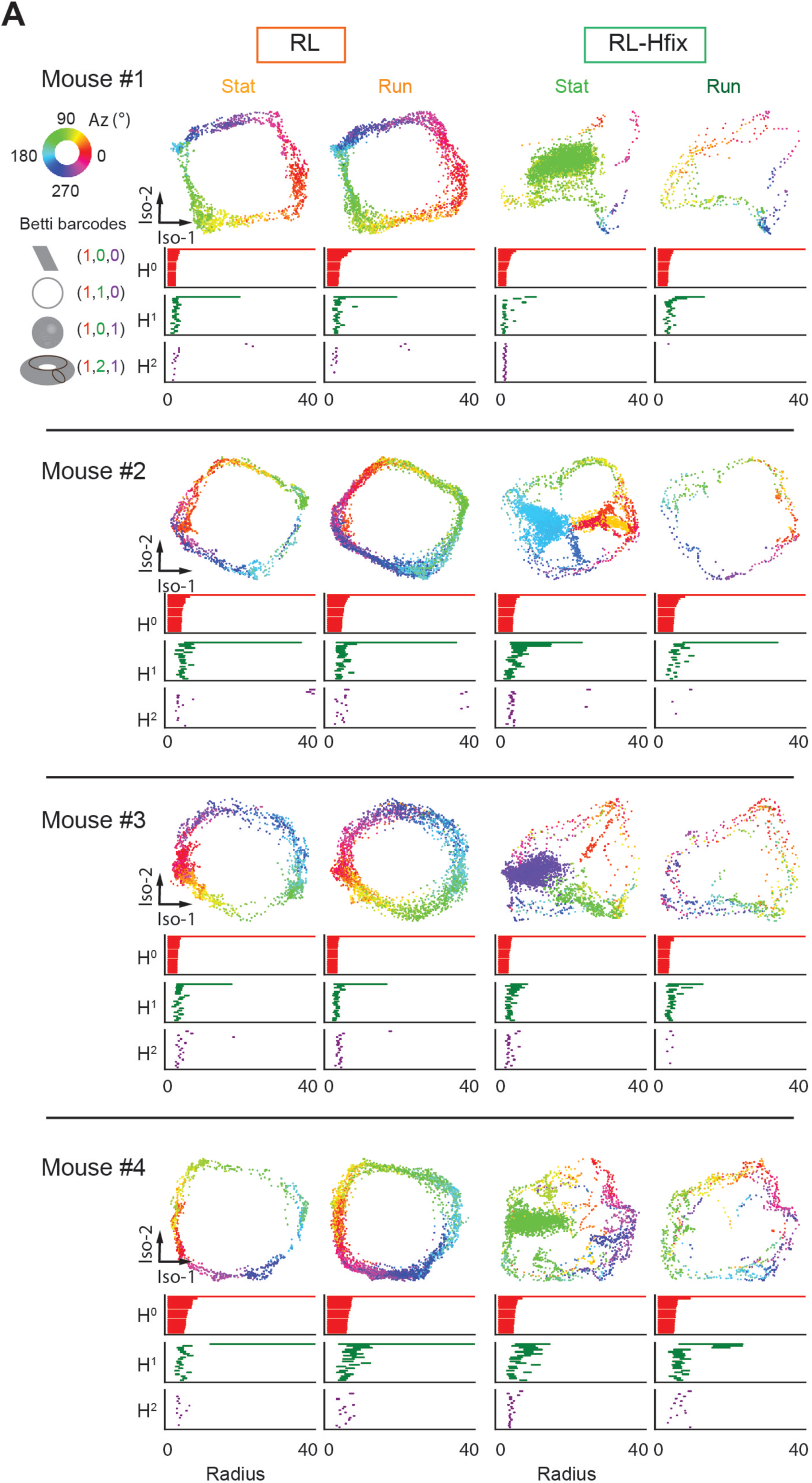
Off-ring dynamics in RL-Hfix across mice. **(A)** Example Isomap projections and corresponding Betti barcodes during Stat and Run epochs for RL and RL-Hfix sessions. Each row represents a different mouse.

**Figure S6:**
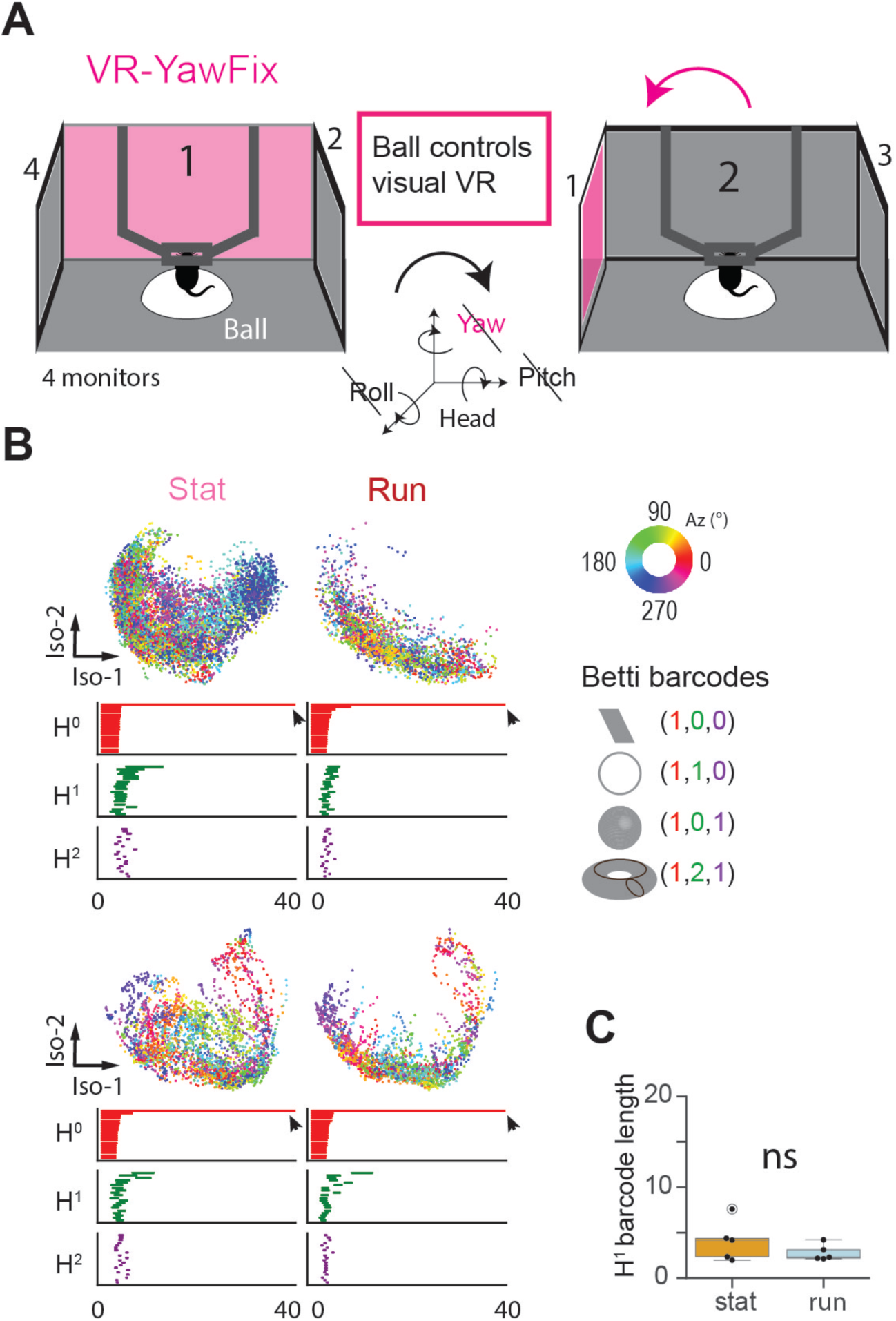
Locomotion fails to restore ring topology in fully head-fixed VR (VR-YawFix). (A) Schematic of VR-YawFix setup. Mice were head-fixed with yaw rotation clamped, preventing angular vestibular input. Animals ran on a spherical treadmill that controlled visual flow in a 2D virtual environment presented on four monitors. Pitch and roll of the head were also fixed. (B) Isomap projections of population activity in VR-YawFix during stationary (Stat, left) and running (Run, right) epochs. Points are colored by decoded azimuth. Corresponding Betti barcode plots show topological features. Loss of the H¹ bar in both states indicates collapse of the canonical HD ring manifold. Two example sessions are shown. (C) No significant difference was observed between states. Wilcoxon signed-rank test H1 bar stat vs run VR-YawFix: median = 4.1 vs 2.3, W = 3.0, P = s0.31, n = 5 sessions).

